# Cryo-EM structure of the *Agrobacterium tumefaciens* T-pilus reveals the importance of positive charges in the lumen

**DOI:** 10.1101/2022.04.28.489814

**Authors:** Jaafar Amro, Corbin Black, Zakaria Jemouai, Nathan Rooney, Caroline Daneault, Natalie Zeytuni, Matthieu Ruiz, Khanh Huy Bui, Christian Baron

**Affiliations:** Department of Biochemistry and Molecular Medicine, Faculty of Medicine, Université de Montréal, Québec, Canada; Department of Anatomy and Cell Biology, Faculty of Medicine and Health Sciences McGill University, Québec, Canada; Montreal Heart Institute, Université de Montréal, Québec, Canada; Department of Nutrition, Faculty of Medicine, Université de Montréal, Québec, Canada

**Keywords:** Type IV secretion system, pili, VirB2, plasmid, conjugation, *Agrobacterium tumefaciens*, pKM101

## Abstract

*Agrobacterium tumefaciens* is a natural genetic engineer that transfers DNA into plants and this is the most frequently applied process for the generation of genetically modified plants. DNA transfer is mediated by a type IV secretion system localized in the cell envelope and extracellular T-pili. We here report the cryo-electron microscopic structures of the T-pilus at 3.2Å resolution and that of the related plasmid pKM101-determined N-pilus at 3Å resolution. Both pili contain a main pilus protein (VirB2 in *A. tumefaciens* and TraM in pKM101) and phospholipids arranged in a 5-start helical assembly. They contain positively charged amino acids in the pilus lumen and the lipids are positively charged in the T-pilus (phosphatidylcholine) conferring overall positive charge to the lumen. Mutagenesis of the lumen-exposed Arg91 residue in VirB2 resulted in protein destabilization and loss of pilus formation. Our results reveal that different phospholipids can be incorporated into type IV secretion system pili and that the charge of the lumen is of functional importance.

## Introduction

The process of DNA transfer from bacteria into other organisms by conjugation is of fundamental and applied scientific importance. Conjugative transfer of DNA is extensively applied in biotechnology for the creation of genetically modified plants using *Agrobacterium tumefaciens* and for the transient expression of proteins for vaccine production (1-3). Simultaneously, bacterial DNA conjugation has significant health implications as plasmid transfer is a major factor contributing to the spread of antibiotic resistance genes (4, 5). Conjugation is mediated by type IV secretion systems (T4SS) that comprise 12 proteins (VirB1 to VirB11 and VirD4) in the archetypical model from *A. tumefaciens* (6-8). The T4SS assembles in the bacterial cell envelope and forms a transmembrane complex and surface-exposed pili in Gram-negative bacteria (9-11). This translocation machinery is highly adaptable to different substrates and it has the capacity to translocate DNA as well as protein substrates across the cell envelope, e.g., single-stranded DNA molecule linked to VirD2 as well as VirE2 in the case of *A. tumefaciens*. The T4SS-determined pilus from *A. tumefaciens* (T-pilus) comprises the main component VirB2 along its entire length and the minor component VirB5 that localizes at the pilus tip (12-14). This primary organization is shared by many T4SS such as the conjugative pili from the IncN plasmid pKM101 (major component TraM, minor component TraC) and even the distantly related T4SS from the human pathogen *Helicobacter pylori*, which carries a tip-localized VirB5 homolog (CagL) (15-17). In contrast, T4SS-determined pili from more distantly related conjugation systems such as plasmids RP4 and F are not known to contain minor components at the pilus tips, but their overall structure is conserved and their main components are homologs of VirB2 (18-20). Although bacterial conjugation has been known for more than 70 years, we are still uncertain whether the T4SS’s substrates (single-stranded DNA and proteins) are actually translocated through the elongated pilus (18, 21-23). Alternatively, this extracellular structure may mediate donor-recipient cell interaction, e.g., via VirB5-homologous tip proteins, followed by pilus retraction, cell fusion and DNA transfer (24, 25).

VirB2 homologs are relatively small proteins comprising 100 to 140 amino acids that often contain long signal peptides. Following their enzymatic processing, the mature 60 to 80 amino acids long pilins are predicted to remain anchored to the inner membrane by two transmembrane helices (20, 26). The biochemistry of the main pilus components VirB2 from *A. tumefaciens* and of its homolog TrbC from the RP4 plasmid has been well characterized and previous data suggested that both undergo specific processing reactions leading to the formation of cyclic peptides (20, 27). However, the cryo-electron microscopy (cryo-EM) analyses of the F-like pilus from the pED208 plasmid and the IncFIIK pilus from the pKpQIL plasmid structures that were previously determined at 3.6Å and 3.9Å, respectively, do not provide any evidence for cyclic peptides (28, 29). Interestingly, their structural analysis revealed the presence of phosphatidylglycerol lipids that were present in a 1:1 stoichiometry in relation to the VirB2-homologous pilin TraA in F-pili. Amino acids from TraA that are exposed to the pilus lumen are overall positively charged, but the presence of phosphatidylglycerol reverses the electrostatic potential to create a negatively charged environment (28, 29). The discovery of these phospholipids may impact pilus function since negatively charged DNA may be transferred through the lumen of the pilus during the conjugation process.

Here, we present the cryo-EM analysis of the *A. tumefaciens* T-pilus and the pKM101 N-pilus at 3.2 and 3Å resolution. Analysis of these structures did not provide evidence for the presence of cyclic peptides, and instead, the T-pilus may contain a novel protein modification at a Cys residue. Intriguingly, we showed that the T-pilus contains the positively charged lipid phosphatidylcholine conferring an overall positive charge to its lumen, whereas the N-pilus contains phosphatidylglycerol. This finding has implications for the passage of substrates across the T-pilus and mutagenesis of positively charged Arg91 of VirB2 demonstrates that its charge is of functional importance.

## Material and Methods

### Bacterial strains and culture conditions

*A. tumefaciens* strain C58 was cultivated on AB minimal medium in the presence of 200 μM of the plant metabolite acetosyringone for virulence gene induction (13, 30) at 20ºC. *E. coli* strain BL21 carrying pKM101 plasmid was cultivated on LB agar with 100 μg/ml ampicillin for plasmid maintenance at 37ºC (12, 14). For large-scale purification, bacteria were cultivated on up to 80 agar plates of 15 cm diameter.

### Tumor formation by *A. tumefaciens*

The wild-type strain C58 and its *virB2* deletion variant CB1002 (31) complemented with plasmids expressing VirB2 or mutant derivatives were applied to wounded leaves of the plant *Kalanchoë diagremontiana*, followed by scoring of tumor formation 2-3 weeks after infection (13).

### Site-directed mutagenesis

The gene encoding VirB2 was mutagenized using QuikChange mutagenesis to change amino acid 91 from Arg to Ala, Glu and Lys and the mutant proteins were expressed in strain CB1002 from plasmid pTrc200 as described (31).

### Expression of VirB proteins and identification of pilins

The levels of VirB proteins (VirB2, VirB5 and VirB8) were analyzed after the separation of samples by SDS-PAGE on 15% Tricine gels (32) and western blotting with specific antibodies as described (13, 14, 33). VirB2 and TraM were identified by silver staining of SDS gels, excised from the gel, followed by proteomics analysis (proteolytic digestion followed by LC-MS/MS) in the institutional service facility at the Institut de recherche en immunologie et cancérologie (IRIC) at Université de Montréal.

### Isolation of pili

The cells were gently scraped off agar plates into 10 mM Na phosphate buffer (pH 5.3) and collected by centrifugation at 15,000x*g*. Pili were released from the cells by passing the bacterial suspension through a hypodermic needle (25 gauge) five times. Next, the cells were removed by centrifugation at 15,000×g, and the supernatant was subjected to centrifugation at 150,000xg to collect the pili fraction. Crude pili (contaminated with bacterial debris and membrane vesicles) were resuspended in 10 mM Tris buffer (pH 7.5, 100 mM NaCl) and 0.5% sodium deoxycholate and further fractionated by velocity sedimentation in a 20-70% linear sucrose gradient at 150,000xg. Fractions containing pili were identified by SDS-PAGE and Western blotting with VirB2- and VirB5-specific antibodies and silver staining, pooled, diluted, and concentrated by centrifugation at 150,000xg to remove the sucrose. Pili were then resuspended in 100 μl of 10 mM Tris buffer (pH 7.5, 100 mM NaCl) followed by visualization by electron microscopy.

### Electron microscopy

For screening of sample quality, 300 mesh copper grids with standard pure carbon film (5 nm) were negatively glow-discharged, and samples were spotted onto the grids and side blotted using Whatman filter paper, followed by staining with 0.75% uranyl formate solution, side blotting and drying at room temperature. The samples were imaged using an FEI Tecnai T12 120 kV TEM (equipped with Gatan 2K AMT and FEI Eagle 4K cameras) in the Université de Montréal EM facility.

Cryo-EM grid preparation was performed using a Vitrobot Mark IV at 4°C and 100% humidity to blot and plunge freeze grids in liquid ethane. For the T-pili, 3 μL of the sample was repeatedly applied and manually blotted three times onto 300 mesh R1.2/R1.3 C-flat Holey carbon grids (Electron Microscopy Sciences, Hatfield, PA, USA) using the multiple blotting technique prior to the Vitrobot step to increase the concentration of pili (34). For the N-pili, the same method as for the T-pili was used, except that C-flat grids were modified to further increase pilus concentration by sputtering 2 nm gold nanoparticles on one side and adding graphene oxide on the other side. The sample was applied on the gold-coated side of the grids before blotting.

Data from vitrified grids were collected by the FEI Titan Krios 300 kV Cryo-STEM equipped with Gatan GIF BioQuantum LS and Gatan K3 Direct Detection camera at the Facility for Electron Microscopy (FEMR) at McGill University. The data were acquired using SerialEM (35) using beam shift with a pattern of 4 holes per movement and 4 movies per hole at 81,000 times magnification and a pixel size of 1.09 Å. The defocus ranged between 0.5 μm to 3.0 μm and movies consisted of 40 frames with a total of 40 e/ Å^2^. A total of 4,224 and 3,744 movies were collected for T-pili and N-pili, respectively.

### Single particle electron microscopy data analysis

Data were processed using CryoSPARC (36). After importing raw movies, patch motion correction, and estimating the contrast transfer function (CTF), particles were manually picked from a subset of the data, extracted and classified for template-based autopicking. Best representative classes were used as templates for filament tracer with a separation distance of 14Å for both T-pili and N-pili. After particle extraction, 2D classification and selection of best classes, 123,100 particles for T-pili and 26,932 particles for N-pili at 256 × 256 box size were selected. Initial volumes without symmetry imposed were constructed using a cylinder as a reference. Using these initial volumes, the helical rise and twist were determined to be 13.832Å and 32.579 degrees for T-pili and 13.839Å and 32.578 degrees for N-pili with 5-start helices configuration. After iterative helical refinement, local and global CTF refinement and symmetry expansion (615,500 particles for T-pili, 134,660 particles for N-pili) were conducted, leading to maps that were refined at 3.2 Å and 3Å for the T- and N-pili, respectively. Initial models were generated automatically using map_to_model job Phenix function (37), then manually fixed in COOT (38) and real space refined using Phenix (37). The pilus assembly was symmetrized using ChimeraX. Visualization was done with Chimera and ChimeraX (39, 40).

For the 3D classification of the T-pili, we first created a mask containing only one lipid and four surrounding VirB2 molecules. Then, we created four references: (1) with full lipid density; (2) with no lipid density; (3-4) with either one of the lipid’s fatty acid tails using the Volume Eraser function of Chimera. For the 3D classification using CryoSPARC, we classified the symmetry expanded particles using “input” mode with four initial references within the mask of one asymmetric unit above to perform a supervised classification to see whether we can extract the heterogeneity of lipids in our data. The distributions of the four classes are 50%, 21%, 12% and 17%, respectively. This indicates that the initial references did not bias the classification. In addition, this distribution is consistent with the heterogeneous nature of the lipid composition revealed by the lipidomic analysis. To further remove the initial model bias, we refined each class with the reference containing the full lipid density. As the resulting cryo-EM map of each class always shows the correct lipid density as indicated in the 3D classification, we conclude that our observation of lipid heterogeneity origin is not a result of any input model’s bias.

### Pilus inner and outer diameter measurements

We first identified the innermost or outermost residues located on the pili (Arg91 and Gly121 for T-pilus, Pro69 and Ser97 for N-pilus, and Asp2 and Thr40 for F-pilus – PDB 5leg). Then, we calculated the X and Y coordinate of the center of the pilus by averaging all the X and Y coordinates of the CA of the residues for the innermost and outermost residues respectively. The inner and outer radii from each residue to the calculated center were calculated and averaged separately. The average and standard deviation for the inner radius is 13.22Å ± 0.02 for T-pilus, 12.00Å ± 0.02 for N-pilus and 15.23Å ± 0.07 for T-pilus. The average and standard deviation for the inner radius is 39.78Å ± 0.02 for T-pilus, 38.81Å ± 0.03 for N-pilus and 43.23Å ± 0.11 for F-pili.

### Pilus alignment

We used ChimeraX matchmaker function to align VirB2, TraM and TraA.

### Interaction analysis

The interaction analysis between different pilins and between pilins and phospholipid was done by PDB sum (41). The interactions between lipid and pilin were visualized with LigPlot+ (42). The surface charges of T-pilus, N-pilus and F-pilus were calculated using ChimeraX.

### Lipidomics analysis

Lipids from pilus samples and bacterial membrane fractions were processed using the liquid chromatography quadrupole time of flight mass spectrometry (LC-QTOF / MS-MS) pipeline at the Montreal Heart Institute as previously described (43). Briefly, pilus samples and bacterial membrane fractions (in 100 μl) were processed after spiking samples with the following six lipid standards: LPC 13:0, PC14:0/14:0, PC 19:0/19:0, PE17:0/17:0, PS12:0/12:0 and PG15:0/15:0. Lipids were extracted into a methyl *tert*-butyl ether (MTBE/methanol/ H_2_0) mixture, followed by further extraction with ethyl acetate, drying under vacuum, and solubilization in methanol/chloroform by vortexing and sonication. The samples (2μl for pili and 1μl for bacterial membrane fractions) were injected into a 1290 Infinity HPLC coupled to a 6550 accurate mass QTOF MS system (Agilent) equipped with a dual electrospray ionization (ESI) source (Agilent) and analyzed in positive scan mode. Lipids were eluted using a Zorbax Eclipse plus C18 reverse-phase column (2.1 × 100 mm, particle size 1.8μm, Agilent) over 83 min at 40ºC using a gradient of solvent A (0.2% formic acid and 10 mM ammonium formate in water) and B (0.2% formic acid and 5 mM ammonium formate in methanol/acetonitrile/MTBE, 55:35:10 (v/v/v)). MS data processing was achieved using Mass Hunter Qualitative Analysis software package (version B.06, Agilent). Finally, lipid annotation, including FA side chains, was achieved by alignment with our in-house database containing more than 500 lipids which have been previously identified in human plasma by MS-MS analysis for which spectra were manually interpreted, similar to Godzien et al. (44).

## Results

### Purification of *Agrobacterium* T-pili and pKM101 N-pili

While the F-like pili can be purified from bacteria cultivated in liquid cultures (28, 29), T-pili and N-pili can be radially detached from their bacterial cells. Accordingly, therefore both the T-pili and N-pili were purified from bacteria grown on semi-solid media. To this effect, *A. tumefaciens* wild-type strain C58 was cultivated on minimal medium agar plates in the presence of the plant metabolite acetosyringone to induce expression of the *virB* genes (13, 14). In contrast, the *tra* genes encoding the pKM101 T4SS are constitutively expressed at a low level (15) and for pili isolation, we cultivated *Escherichia coli* strain BL21 carrying the plasmid on LB agar plates. The pili were purified by shearing concentrated bacteria samples through hypodermic needles, differential centrifugation to separate the cells, ultracentrifugation to enrich the pili and finally, sucrose gradient sedimentation (Fig. 1). In the case of *A. tumefaciens*, the different purification steps were monitored by Western blotting with VirB2- and VirB5-specific antisera and silver staining revealing increasing purity and mass spectrometry confirmed the presence of VirB2 in the purified samples (Fig. 1A and supplementary Fig. 1). Surprisingly, the signal for the tip-localized VirB5 was lost after the initial purification steps, indicating that the tip protein may be easily broken off and/or detached by the addition of sodium desoxycholate required for optimal purification of the sample. IncN pili were purified from *E. coli* using the same procedure, but due to the lower expression of these pili, the final sample was not as pure as the T-pili sample. Since we have no specific antisera developed against the main pilus component TraM, the purification was monitored by silver staining and the identity of the pilin in the final sample was confirmed by mass spectrometry (Fig. 1B and supplementary Fig. 1). The purified pili were next analyzed by negative staining and transmission electron microscopy (TEM), showing the presence of ∼10 nm diameter T-pili (Fig. 1C) as well as ∼10 nm diameter N-pili (Fig. 1D).

**Figure 1.**
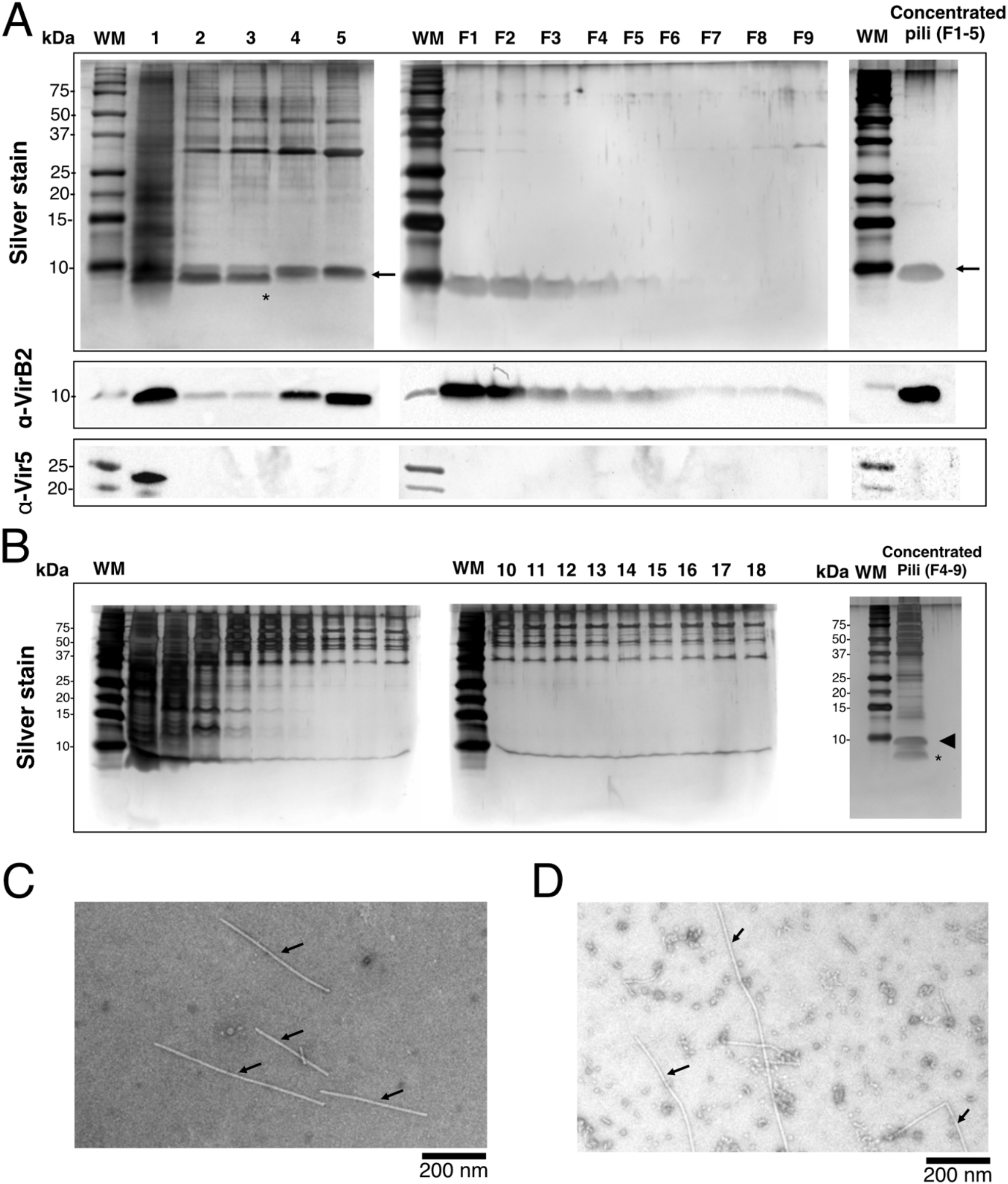
Purification of T-pili from *A. tumefaciens* and of pKM101 N-pili. (A) Purification of T pili through successive steps of sodium deoxycholate treatment and sucrose density gradient centrifugation as described in materials and methods. Samples were resolved by tricine-SDS-PAGE and analyzed by silver staining (*top panel*) and western blotting using VirB2 (*middle panel*) or VirB5 (*lower panel*) specific antiserum. (*1)* Crude extracted pili containing flagella, membrane vesicles and membrane debris, (*2-5*) crude pili after treatment 3 times with 0.5% sodium deoxycholate to remove contaminants. (*F1 to F9*) Fractions from the top of the 20-70% sucrose gradient. (WM) weight marker. The numbers on the left indicate the molecular weight in kDa. Arrows indicate VirB2. Asterisk indicates a contaminant, probably the major outer membrane lipoprotein, removed by the sodium deoxycholate detergent. VirB2 identity confirmed by mass spectrometry analysis. (B) Tricine-SDS-PAGE and silver staining of 18 fractions from the top of the 20-70% sucrose gradient of crude extracted N-pili. Fractions were analyzed by electron microscopy and N-pili were detected abundantly in fractions 4 to 9, which were then pooled and concentrated. Arrow’s head indicates TraM. Asterisk indicates the major outer membrane lipoprotein LPP. The identities of TraM and LPP were revealed by mass spectrometry analysis. T-pili (C) and N-pili (D) visualized by electron microscopy. Scale bar 400 nm. Arrows indicate pili.

### Cryo-EM analysis reveals that T-pili and N-pili comprise 5-start helical assemblies

The quality of the samples comprising T- and N-pili was next assessed by cryo-EM on a lower resolution 100 kV instrument and whereas their concentrations were not high (2-3 pili per hole for the T-pilus and 4-5 pili per hole for N-pili), they readily entered the holes of the EM grids (supplementary Fig. 2A, B) enabling subsequent data collection on a 300 kV Titan Krios instrument. Data were processed using CryoSPARC, which resulted in 3.2 Å and 3Å resolution maps for the T- and N-pili, respectively (supplementary Fig. 2C, D, E). The overall dimensions of T-pili (outer diameter of 79.6Å, lumen diameter of 26.4Å; Fig. 2A, B) and N-pili (outer diameter of 77.6Å, lumen diameter of 24.0Å; Fig. 2F, G) are similar. The major pilus components VirB2 (Fig. 2C) and TraM (Fig. 2H) are arranged in 5-start helical assemblies and the helical rise and helical twist are comparable (13.863Å and 32.579° for T-pilus; 14.708 Å and 30.235° for N-pilus) (Fig. 2E, I). Analysis of the VirB2-VirB2 interactions in the T-pilus and TraM-TraM interactions in the N-pilus revealed that each pilin subunit forms extensive interactions with six other subunits (areas of interaction interface shown in supplementary Fig. 3A, B, C) primarily via hydrophobic interactions (supplementary Fig. 3D, E). The four N-terminal amino acids (Q/S/A/G) of VirB2 (Fig. 2C) are not modeled due to flexibility. In contradiction with the previous proteomics analysis of VirB2 (20), both proteins do not form cyclic peptides. Interestingly, we observed an extended electron density on amino acid Cys64 of VirB2 (Fig. 2J) that could not be resolved at the current resolution and proteomics analysis did not reveal the identity of the modification either. We did not observe a comparable extended density on the corresponding amino acid Gln41 of TraM (Fig. 2K).

**Figure 2.**
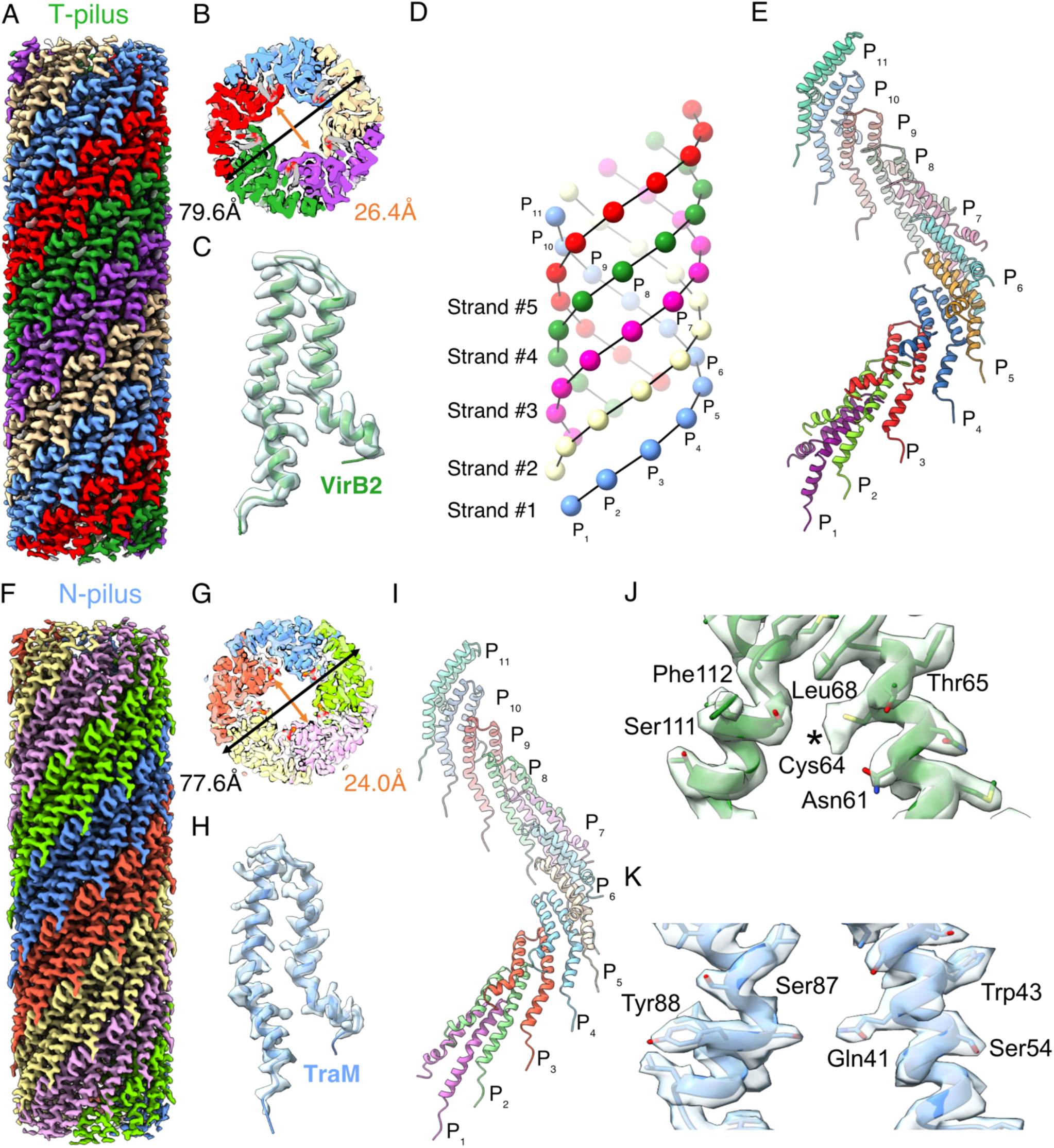
Cryo-EM maps of T-pili and N-pili. Cryo-EM map of T-pilus (A) and N-pilus (F) showing the 5-start helix indicated by different colors. Cross-section of the T-pilus (B) and N-pilus (G) showing the measured inner and outer diameters. The density and models of a single unit of VirB2 (C) and TraM (H). (D) A cartoon showing a complete repeating unit and the naming system of the T-pilus with 5 strands (Strand #1-5) and each strand consists of 11 pilins (P_1_ to P_11_). A single twist of 11 units from T-pilus (E) and N-pilus (I). (J) Cys64 of T-pilin showing a bulky density (noted with asterisk *) does not correspond to the cysteine side chain, which might correspond to a novel modification. (K) The Gln41 of TraM does not show a similar additional density as Cys64.

### The *A. tumefaciens* T-pilus contains phosphatidylcholine lipids

Lipids are present in a one-to-one stoichiometry to the TraA pilin of the F-pilus (28) and we observed similar amounts of phospholipids in the T-pilus (Fig. 3A, B) and the N-pilus (supplementary Fig. 4A, B). The electron density of the lipid in the N-pilus is similar to phosphatidylglycerol in the F-pilus (Fig. 3C, D), but the lipid in the T-pilus has an extended head group that could be modeled to fit phosphatidylcholine (Fig. 3C). To identify this lipid, we subjected cell membranes and isolated T-pili to lipidomics analysis revealing the predominant presence of diacyl-as well as monoacyl-phosphatidylcholine in the membranes and the isolated T-pili (Table 1). The phospholipids are deeply embedded in the helical assembly (Fig. 3E, F for T-pilus and supplementary Fig. 4C for N-pilus), and the acyl groups of each molecule interacts with five pilins primarily via hydrophobic interactions (Fig. 3F and supplementary Fig. 4C, E, F). In addition, the choline head group is exposed to the lumen of the pilus (Fig. 3B), contributing to its overall positive charge (see below).

**Figure 3.**
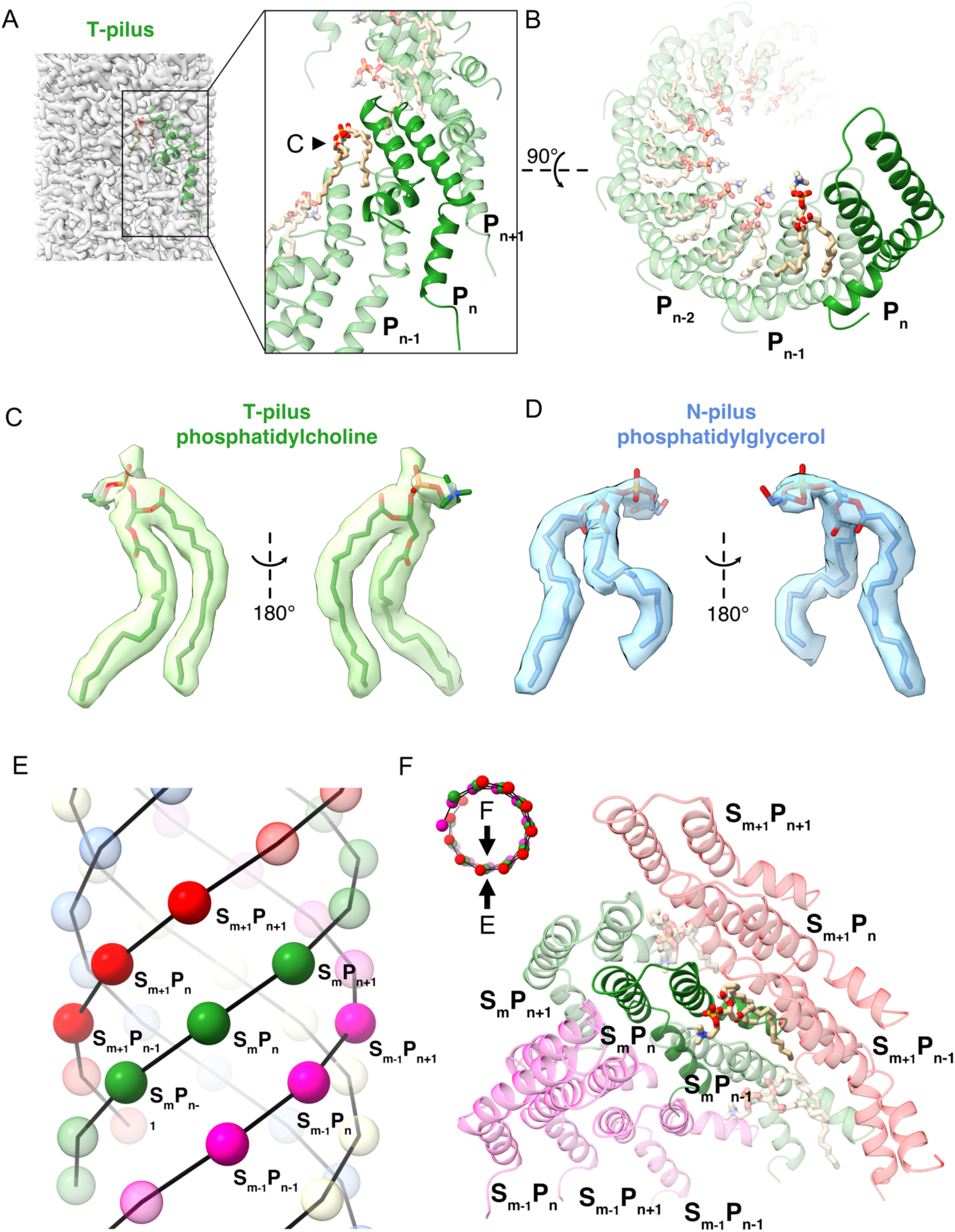
T-pili contain phospholipids. (A) T-pili contain phospholipids in a stoichiometry of 1:1 with VirB2. Diacyl-phosphatidylcholine was modelled into the density. The arrow indicates the view of the phosphatidylcholine in (C). (B) Arrangement of VirB2 pilin and phosphatidylcholine in a single helical strand. (C) The phosphatidylcholine inside its corresponding density inside the T-pilus. (D) The phosphatidylglycerol inside its corresponding density inside the N-pilus. (E) Schematic cartoon of the T-pilus pilin arrangement showing an interaction unit of nine pilins from three adjacent helical strands. S_m_ denotes Strand #m; P_n_ denotes Pilin #n within a strand. (F) Interaction of VirB2 pilin and phosphatidylcholine within a nine-pilin unit shown in (E). The small cartoon showing the view in (F) indicated by the black arrow.

**Table 1.** Lipidomics analysis of purified T-pili and membranes. Mass spectrometry analysis of the lipids extracted from the T-pilus and *A. tumefaciens* membranes. (A) Comparison of identified lipid entities extracted from the T-pilus, and membranes from acetosyringone-induced (AS) and non-induced *A. tumefaciens* (n = 1). (B) Normalized area under the curve (AUC) values of specifically identified lipids extracted from the T-pilus, reflecting the signal intensity and relative abundance of each lipid (n = 3) that were annotated using the in-house database previously validated by MS/MS analysis. FA: fatty acid, LPC: monoacylglycerophosphocholine, PC: diacylglycerophosphocholine, LPE: monoacylglyceroethanolamine, PE: diacylglyceroethanolamine, CE: steryl ester, DG: diacylglycerol, TG: triacylglycerol

| Sample | Numbers of entities |  |  |  |  |  |  |  |
| --- | --- | --- | --- | --- | --- | --- | --- | --- |
|  | FA | LPC | PC | LPE | PE | CE | DG | TG |
| membranes - AS | 2 | 9 | 28 | 2 | 4 | 1 | 6 | 2 |
| membranes +AS | 2 | 9 | 28 | 2 | 3 | 1 | 6 | 2 |
| T-pili | 2 | 7 | 7 | 0 | 1 | 0 | 4 | 2 |

B
| Lipid ID | AUC* |  |  |
| --- | --- | --- | --- |
|  | Sample #1 | Sample #2 | Sample #3 |
| LPC14:0-b [M+H]<sup>+</sup> | 1123427 | 2011779 | 2210257 |
| LPC19:0-a [M+H]<sup>+</sup> | 495260 | 373694 | 456915 |
| LPC19:0-b [M+H]<sup>+</sup> | 455270 | 385926 | 361127 |
| PC(36:0) [M+H]<sup>+</sup> | 460600 | 375440 | 312769 |
| PC(19:1_18:1) [M+H]<sup>+</sup> | 96520 | 146187 | 758474 |
| DG(16:0/18:0) [M+NH<sub>4</sub>]<sup>+</sup> | 304168 | 304968 | 379923 |
| PC(40:0) [M+H]<sup>+</sup> | 345379 | 248781 | 224882 |
| LPC13:0-a [M+H]<sup>+</sup> | 179411 | 165675 | 175034 |
| PC(18:1_18:1) [M+H]<sup>+</sup> | 52837 | 131464 | 295057 |
| DG(18:0/18:0) [M-H<sub>2</sub>O]<sup>+</sup> | 110696 | 182762 | 166442 |
| DG(16:0/18:0) [M-H<sub>2</sub>O]<sup>+</sup> | 129231 | 104838 | 153003 |

The lipidomic analysis revealed that the lipids we succeeded to annotate in the T-pili are heterogeneous and composed primarily of monoacyl- and diacyl-phosphatidylcholine (Table 1), which does not reflect in our consensus cryo-EM map. To probe the heterogeneity of lipid in T-pili, we attempted to perform a 3D classification at the lipid-containing pocket (Fig. 4A-C). Since unsupervised classification is not sensitive enough to reveal the small difference in lipid structure, we performed a supervised classification based on our knowledge from the lipidomic analysis (Materials & Methods). The symmetry expanded particles are classified into four input references corresponding to diacyl-phosphatidylcholine, no lipid density and monoacyl-phosphatidyl choline-like densities by deleting either fatty acid tails from the original diacyl-phosphatidylcholine density. The 3D classification revealed the heterogeneous nature of the lipids within the lipid pocket. While 50% of the particles contain diacyl-phosphatidylcholine density, 21% do not contain any lipid and 29% present monoacyl-phosphatidylcholine-like densities (Fig. 4D). In addition, the monoacyl-phosphatidyl choline-like densities in class 3 and 4 look larger than expected, indicating a more flexible structure when only monoacyl-phosphatidyl choline is present.

**Figure 4.**
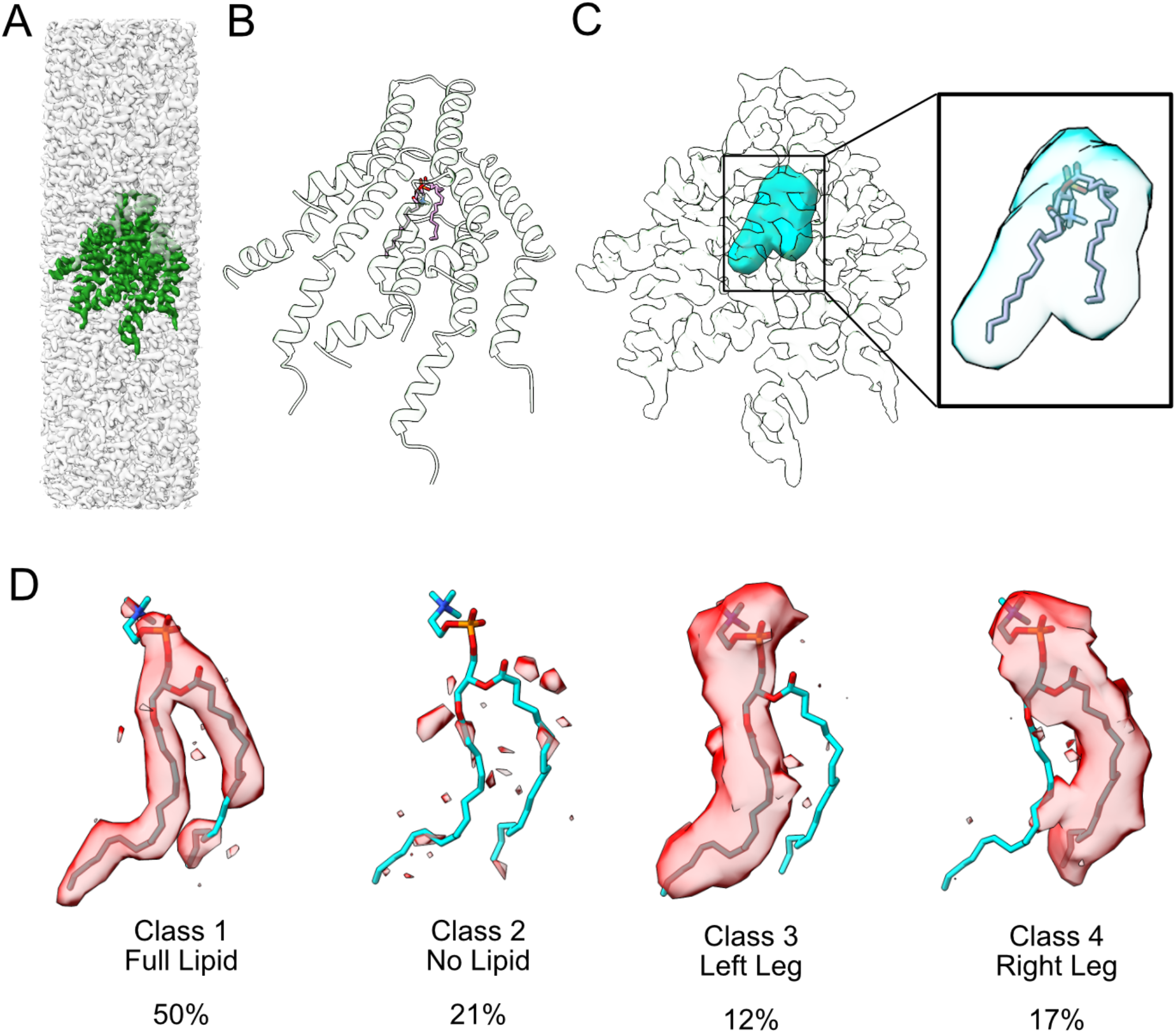
3D classification reveals heterogeneous lipid densities in the T-pili. (A) The consensus T-pili 3D map showing the region subjected to focused 3D classification. (B) The focused classified region contains 1 lipid molecule and 4 VirB2 molecules. (C) Highlight of the lipid pocket (cyan) where the lipid molecule is located in the focused classified region. (D) Zooming into the lipid pocket of 4 classes from 3D classification. Class 1 shows a phosphatidylcholine density. Class 2 shows no lipid density. Class 3 and 4 show lipid density in only the left leg or right leg, respectively; this likely to corresponds to LPC as detected by lipidomics analysis. The thresholds used for all 4 classes are calibrated to make the VirB2 density the same. Therefore, the large densities in the lipid pockets demonstrate that the lipids in Class 3 and 4 are more flexible than that of Class 1.

### Comparative analysis of T-, N-, and F-pili

The availability of two novel and distinct structures of T4SS-determined pili enabled a comparative analysis with the pED208-encoded F-pilus (28). The overall dimensions of the three pili are similar even if the width and the lumen of the F-pilus (external/internal diameter: 86.5Å / 30.5Å) are larger than those of the T- (79.6Å / 26.4Å) and N-pilus (77.6Å / 24.0Å) (Fig. 5A). Despite the low primary sequence identities (9-19%) between the main components of the three pili, their overall structures and fold are very similar (Fig. 5B-D). The matured and processed version of the main components in all three pili; VirB2 (74 amino acids), TraM (70 amino acids) and TraA (64 amino acids), are all short proteins that comprise two hydrophobic α-helices and a loop region that is exposed to the lumen of the pilus that contains positively charged Arg, Lys or His residues (Fig. 5B). VirB2 and TraM are more structurally similar than TraA (Fig. 5C). Despite their structural similarity, the pilins contain differently charged phospholipids with distinct conformations (Fig. 5B) that are likely to be associated with the primary sequence alternations and may impact their function.

**Figure 5.**
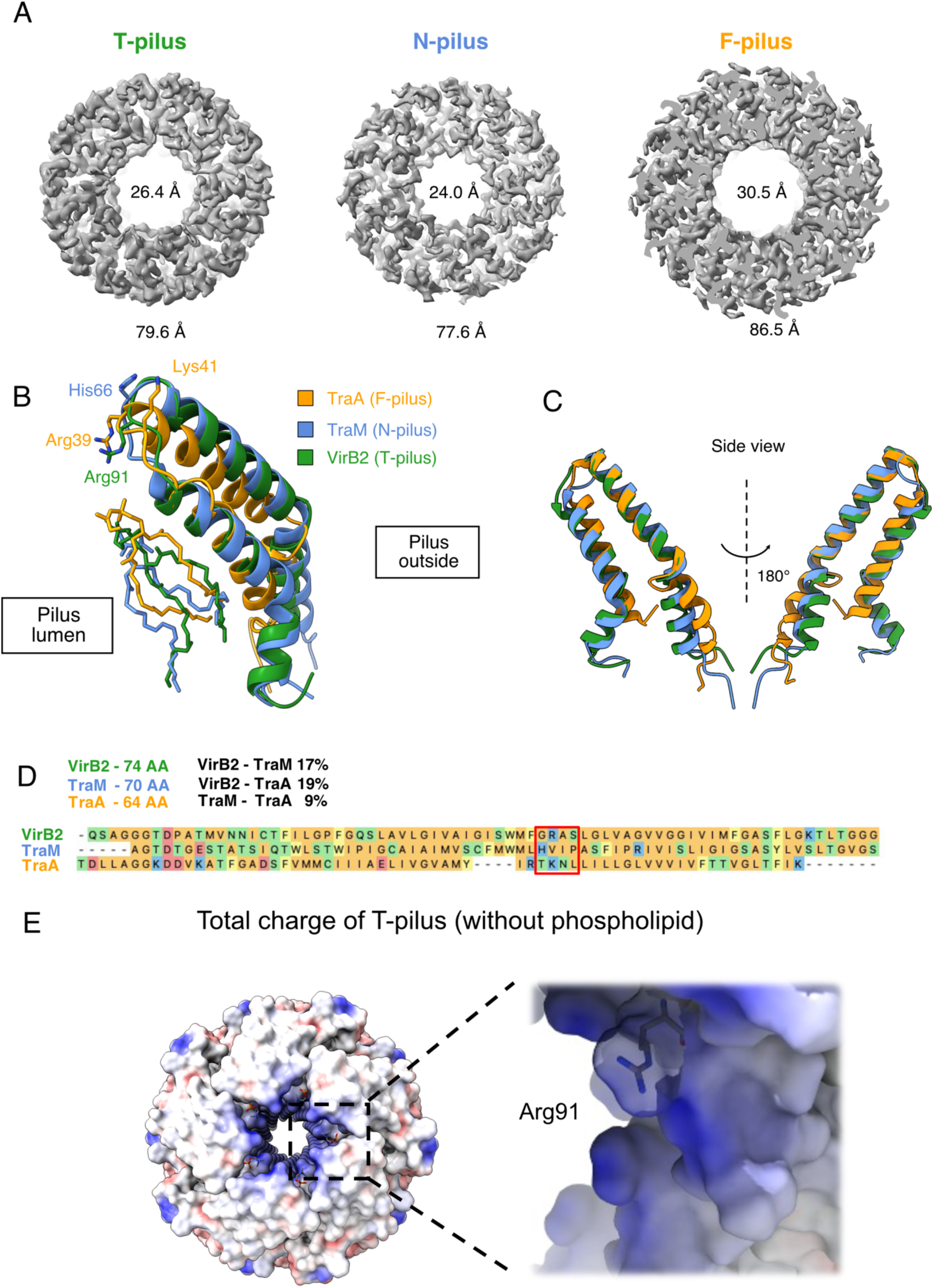
Comparative analysis of T-, N- and F-pili. (A) The F-pilus has a significantly larger lumen diameter (30.5Å) than the T-pilus (26.4Å) and N-pilus (24Å). (B) Overlay of VirB2 (T-pilus), TraM (N-pilus) and TraA (F-pilus) using ChimeraX matchmaker function. The positive residues Arg91 (VirB2), His66 (TraM), Arg39 and Lys41 (TraA), which point into the pilus lumen are shown. The overlay of the lipids of VirB2, TraM and TraA indicates that the lipid position can be flexible relative to the pilin. (C) Side view & top view of TraM, TraA and VirB2 indicate that VirB2 and TraM are more structurally conserved than TraA. (D) The alignment of pilin sequences was generated with MegAlign Pro Software, version 17.2 (DNA Star). (E) The surface charge of the lumen of T-pilus (calculated without the phosphatidylcholines) showing Arg91 pointing toward the lumen and contribute significantly to the positive charge of the lumen.

The F-pilus (28) and the N-pilus (Fig. 5B) contain phosphatidylglycerol and their lumen is overall negatively charged (supplementary Fig 6. A-F). In contrast, the lumen of the T-pilus is overall positively charged and the presence of the lumen-exposed choline group of the phospholipid further increases the positive charge (Fig. 5E). In addition, the outside surfaces of T-pilus, N-pilus and F-pilus display different charge patterns (supplementary Fig. 6A-C). The T-pilus is mostly positively charged while the N-pilus and F-pilus show patches of positive and negative charges.

### The positive charge of Arg91 is important for the functionality of the *A. tumefaciens* T-pilus

To assess the importance of positive charges in the lumen of the T-pilus, we changed the lumen-exposed amino acid Arg91 of VirB2 to Ala (no charge), Glu (negative charge) and Lys (reduced positive charge). The VirB2 wild type and mutant proteins were expressed in *virB2* deletion strain CB1002, followed by virulence gene induction and isolation of T-pili. In parallel, we also infected *Kalanchoë diagremontiana* plants to assess functional complementation. Whole-cell lysates and pilus fractions were analyzed by SDS-PAGE and western blotting with VirB2-specific antiserum revealing the presence of the main pilus component in cells and pilus fractions from C58 wild type *Agrobacterium* and CB1002 complemented with VirB2 (Fig. 6A). In contrast, similar to a previous report (26) after complementation of CB1002 with the VirB2R91A mutant the protein was neither detected in cell lysates nor in pilus fractions. Reduced amounts of the VirB2R91K and of the VirB2R91E mutants were detected in cell lysates of complemented CB1002, but these proteins were not detected in pilus fractions. Consistent with these observations, only expression of VirB2 wild-type protein led to functional complementation (tumor formation) after infection of *K. diagremontiana* plants with CB1002 (Fig. 6B), underlining the importance of positively charged amino acid Arg91 for pilus assembly and function.

**Figure 6.**
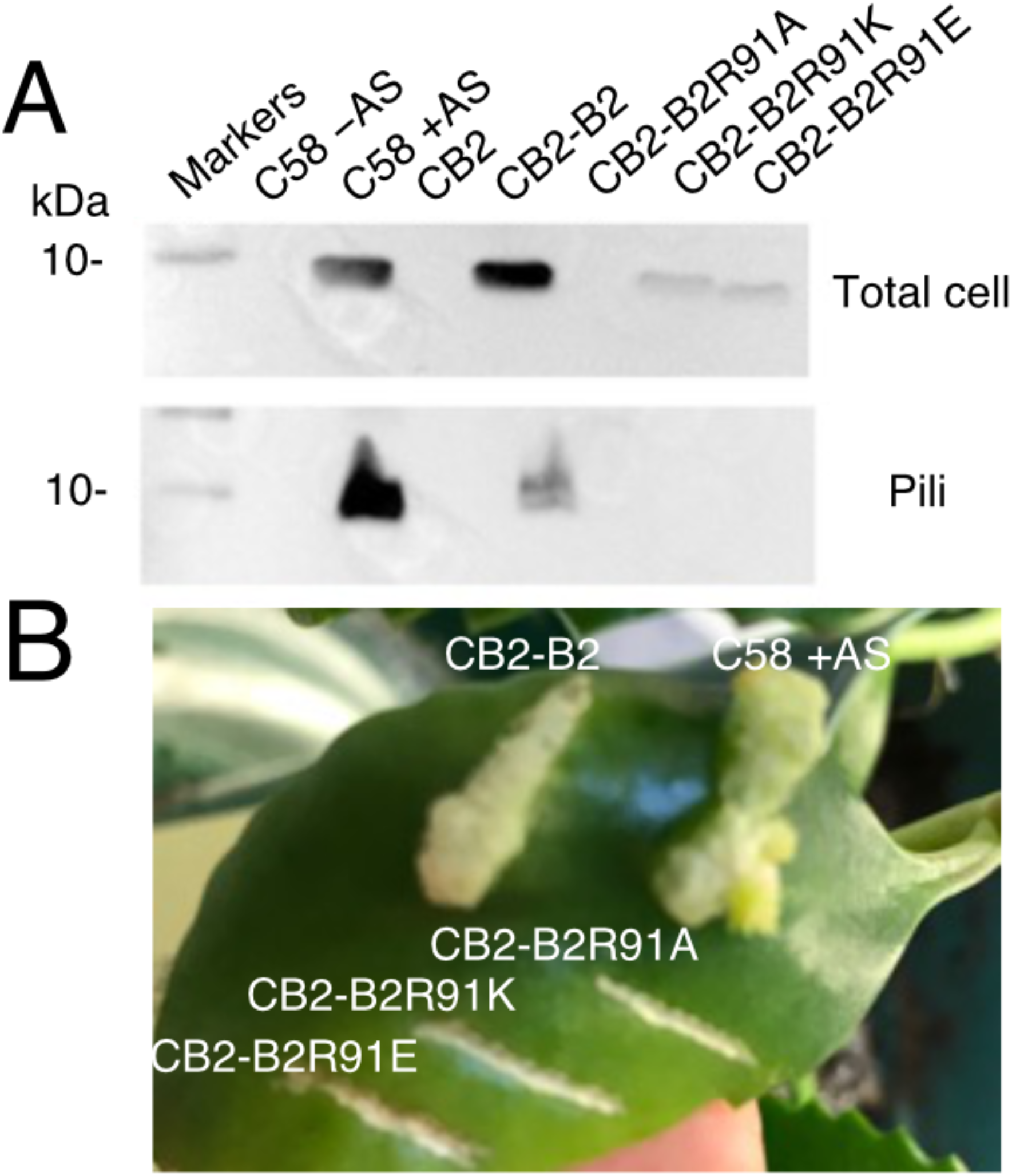
Effects of changes of VirB2 amino acid Arg91. (A) SDS-PAGE and western blotting of cell lysates and isolated pilus fractions with VirB2-specific antisera in C58, CB2 and complemented strains with R91A, R91K and R91E mutants. (B) Tumor formation test after infection of *K. diagremontiana* plant. C58: wild type; AS: acetosyringone for induction of virulence genes; CB2: CB1002, *virB2* deletion strain; CB2-B2: CB1002 strain complemented with VirB2; CB2-B2R91A, CB2-B2R91K, CB2-B2R91E: CB1002 strains complemented with R91A, R91K and R91E mutant VirB2, respectively.

## Discussion

The discovery of abundant amounts of lipids in the F-pilus was a groundbreaking discovery changing the way we contemplate the biology of this cellular appendix. Since its discovery more than 70 years ago, it has been suspected that DNA may be transferred between cells through an extended pilus (45). However, there is still no direct evidence for this hypothesis. According to an alternative model, the pilus may primarily mediate cell-cell contact formation, followed by DNA transfer between closely juxtaposed donor and recipient cells as observed by EM (22, 23) and fluorescence microscopy (24, 25). The present study does not directly address this substrate transfer question but rather supplements the growing body of evidence indicating a significant amount of phospholipid presence in T4SS-determined pili (28, 29). Considering that phospholipids are present in a one-to-one stoichiometry of the pilins, the pilus could be considered as an extension of the bacterial membrane that contributes to cell-cell fusion, followed by the translocation of DNA and other substrates across the T4SS that may or may not include a shortened pilus channel during the transfer process.

The contribution of the lipids to pilus function is still unclear and in previous publications, it was speculated that their negative charge contributing to a negatively charged lumen of the pilus may repel negatively charged DNA and thereby contribute to its transfer (28, 29). This hypothesis was based on the presence of phosphatidylglycerol in F-like pili and this lipid is likely also present in the N-pili described in this work. However, the T-pili from *A. tumefaciens* contains phosphatidylcholine and its lumen is overall positively charged, which is not consistent with a model implying that DNA is being repelled by the negatively charged lumen of the pilus. We mutated Arg91, which is also exposed to the lumen of the pilus, to test the importance of positive charges, and surprisingly, all the changes (Arg91 to Ala, Glu or Lys) reduced the stability of the pilin in the cell and abrogated pilus formation. In contrast, the change of Arg94 (Arg39 of the processed pilin) to Glu did not impact pilus assembly and conjugation of pED208, but it negatively affected phage infection contributing to the notion that the assembled pilus may serve as a conduit for naked DNA (28). Arg94 is not conserved in other F-like pili, but some changes of closely juxtaposed Lys residues that are probably exposed to the lumen of these pili reduced phage infection and conjugation efficiency, but not pilus assembly (46). These results demonstrate that Arg91 and its charge are crucial for VirB2 pilin stability and function, which is quite distinct from the effects of similar changes in F-pili. It will be interesting to assess in the future whether this difference is due to the difference in lipid content (phosphatidylglycerol vs. phosphatidylcholine) and whether the incorporation of these lipids is specific or a diffusion-mediated process driven by the composition of the bacterial membrane.

The discovery of cyclic pilins in VirB2 and its homolog TraC from the RP4 plasmid were intriguing and we have no straightforward explanation for the difference in our results obtained by cryo-EM from previous proteomics-based approaches. Purified *A. tumefaciens* and RP4 pilins were digested with proteases, followed by sequencing and the absolute molecular mass of the processed pilin lacked 18 Da, which was attributed to the formation of an N- to C-terminal peptide bond (20, 27). We did not observe densities for the N-terminal amino acids in VirB2 (QSAG), but due to the distance between the terminal amino acids, it was not possible to model these into a cyclic structure. Nevertheless, our results are consistent with the results of cryo-EM analyses of F-like pili and cyclic versions of the TraA pilins were not observed in both cases (28, 29). The additional density observed on Cys64 of VirB2 may be key to explaining this discrepancy. In principle, this amino acid could form a thioether link with Ser111, thereby forming a cyclic structure and this possibility would be consistent with the results of previous studies (20, 27). However, Cys64 is not essential and changes of this amino acid to Ser of Ala only slightly reduced bacterial virulence at elevated temperatures (26, 47). Therefore, cyclisation may not be essential for the assembly and function of T4SS-determined pili.

Pioneering work has been published on F-pili that are actually are quite distinct from T-pili and N-pili described here (28, 29). Similarly, there are large differences in the protein composition of the T4SS that are much for complex in the case of F-pili (48). F-like pili are often N-terminally acetylated, but this modification is not required for pilus function (46). In contrast, the tip-localized VirB5/TraC proteins are essential components of the *Agrobacterium* and pKM101 secretion systems and they are believed to be involved in contact formation with host cells. In addition, the width and lumen of F-pili are significantly wider than that of T- and N-pili and amino acid changes in the lumen have very different effects on pilus function. Finally, the phospholipids in the T-pili confer a positive charge to the lumen of the pilus and our results suggest the presence of di-acetylated as well as mono-acetylated entities. These differences suggest that there may be significant functional differences between F-like and T-/N-like pili that warrant future studies aimed at elucidating the molecular details of the mechanisms of these fascinating and adaptable macromolecular conduits.

## Acknowledgments

This work was supported by grants from the Natural Sciences and Engineering Research Council (NSERC, #RGPIN-2017-05123) and the Canadian Institutes of Health Research (CIHR, #398288 and #274108) to C.B., funding from FRQNT (Établissement de la relève professorale, #297339) to N.Z., funding from CIHR (PJT-156354) and NSERC (#RGPIN-2016-04954) to H.B. We grateful to staff at the FEMR at McGill, Dr. Kaustuv Basu and Kelly Sears and at the Université de Montréal EM facility for technical support and assistance.

## Author contributions

J.A. designed and carried out experiments and analyzed data; C.B. analyzed data; Z.J. designed and carried out experiments and analyzed data; N.R. carried out experiments and analyzed data; C.D. carried out experiments and analyzed data; N.Z. analyzed data; M.R. designed experiments and analyzed data; H.B. designed experiments and analyzed data; C.B. designed experiments, analyzed data and wrote the manuscript with input from all the co-authors. All co-authors have approved the final version of this manuscript.

## Data Availability

The data produced in this study are available in the following databases:

- Cryo-EM maps of T-pili and N-pili: EMDB EMD-XXXXX and EMD-XXXXX (https://www.emdataresource.org/EMD-XXXXX)
- Model coordinates of T-pili and N-pili: PDB YYYY and YYYY (https://www.rcsb.org/structure/YYYY)

## Supplementary data

**Supplementary table 1.**
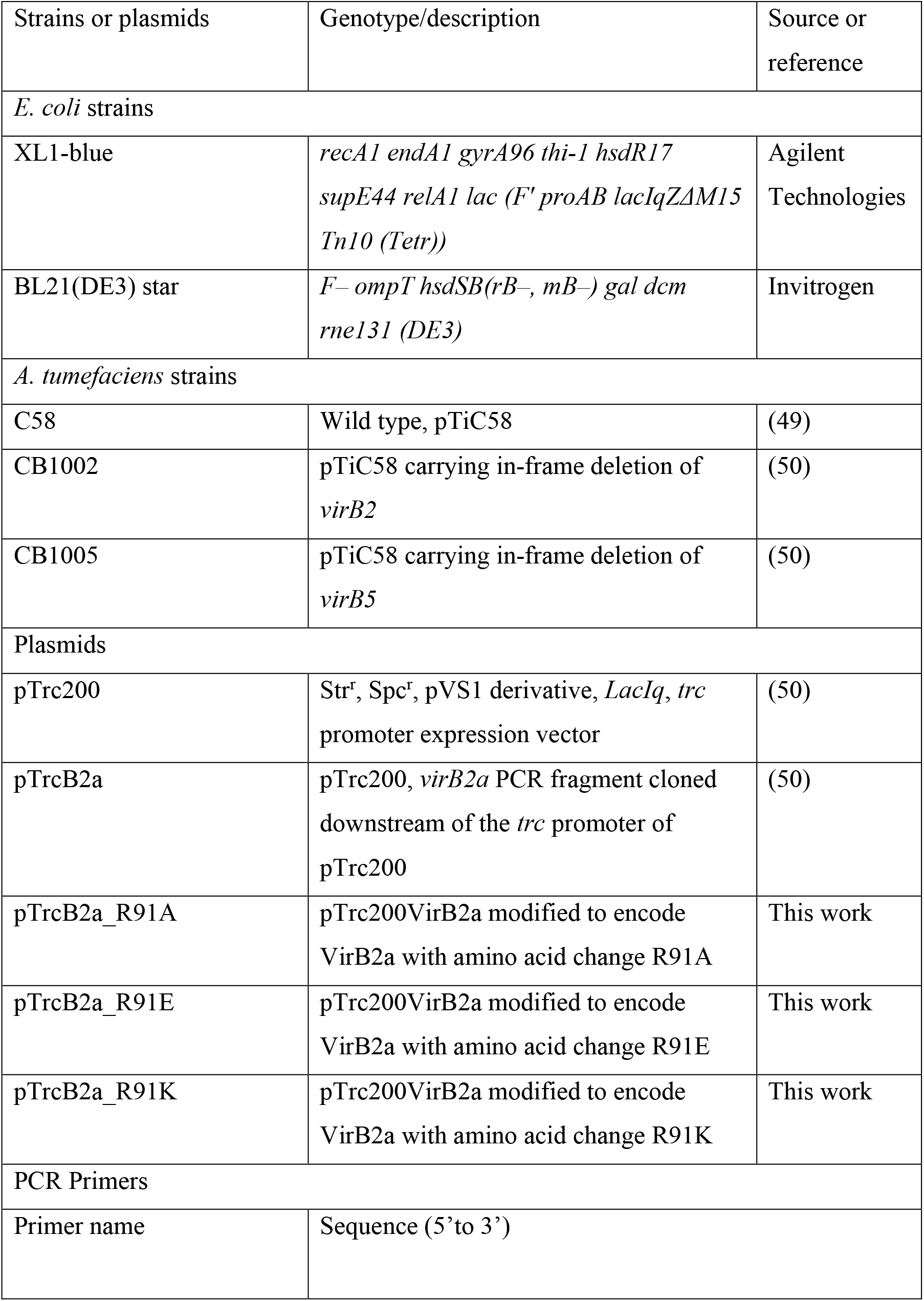
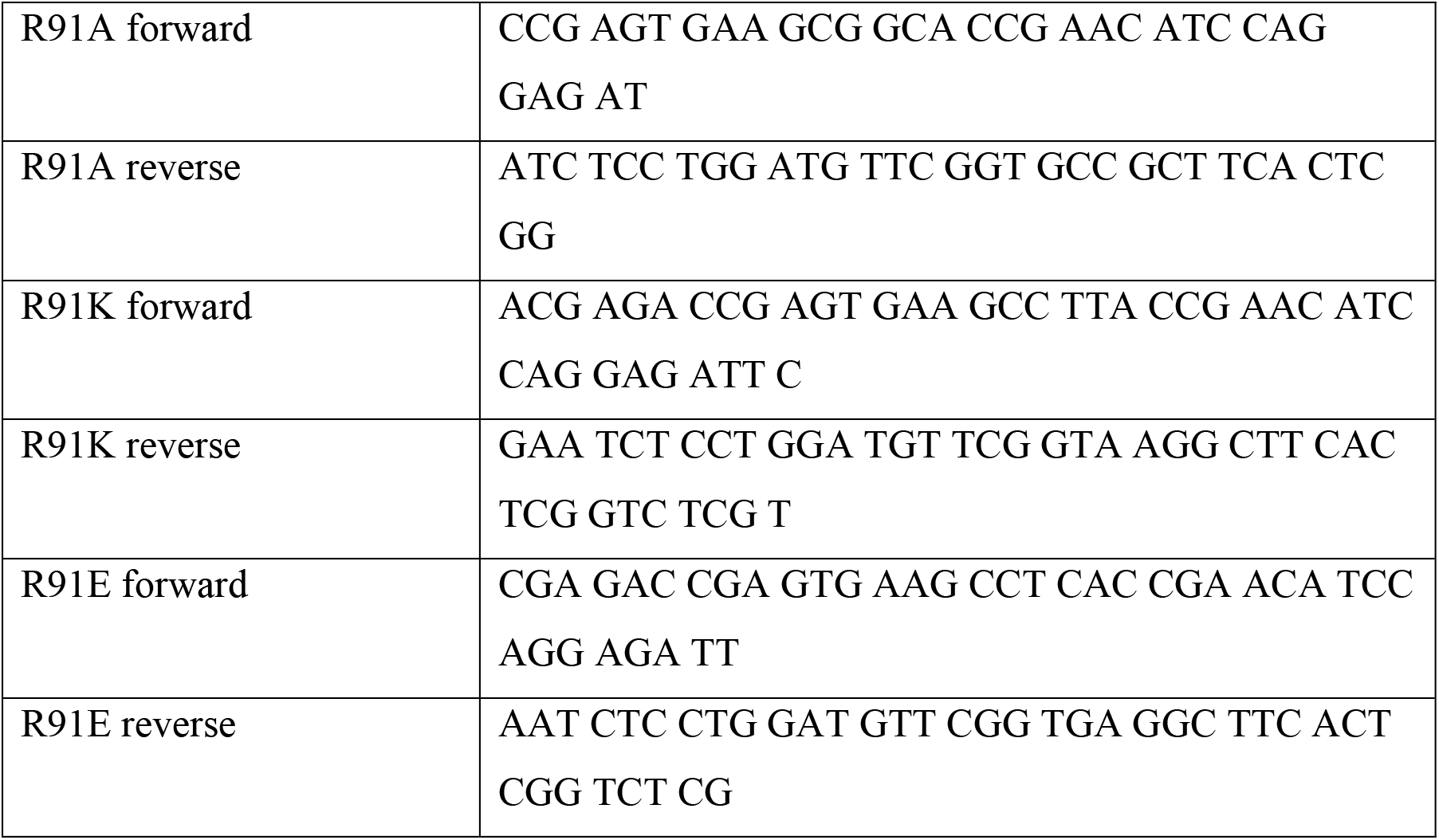
Strains, plasmids and oligonucleotides used for mutagenesis of VirB2.

## Supplementary Figures

**Supplementary Figure 1.**
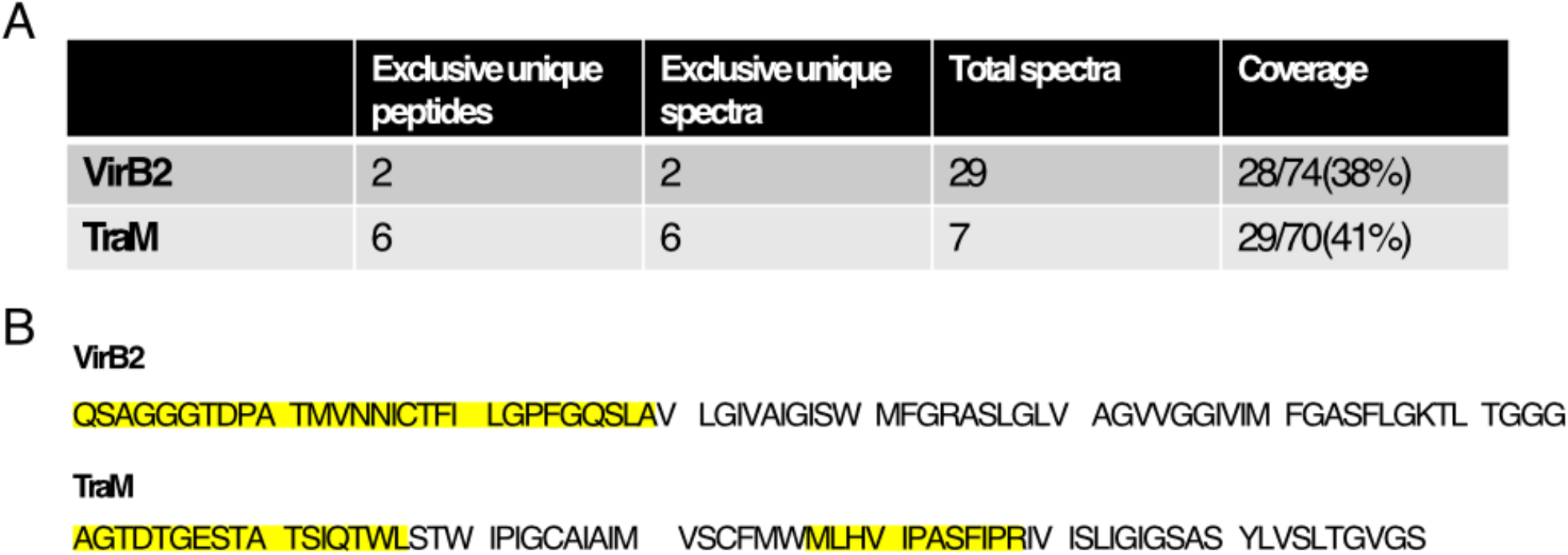
Identification of VirB2 and TraM by mass spectrometry (MS). (A) MS analysis of the ∼10 kDa protein excised from tricine-SDS-PAGE of concentrated T-pilus and N-pilus samples as shown in Fig. 1A and Fig. 1B (B) Identified peptides highlighted in yellow.

**Supplementary Figure 2.**
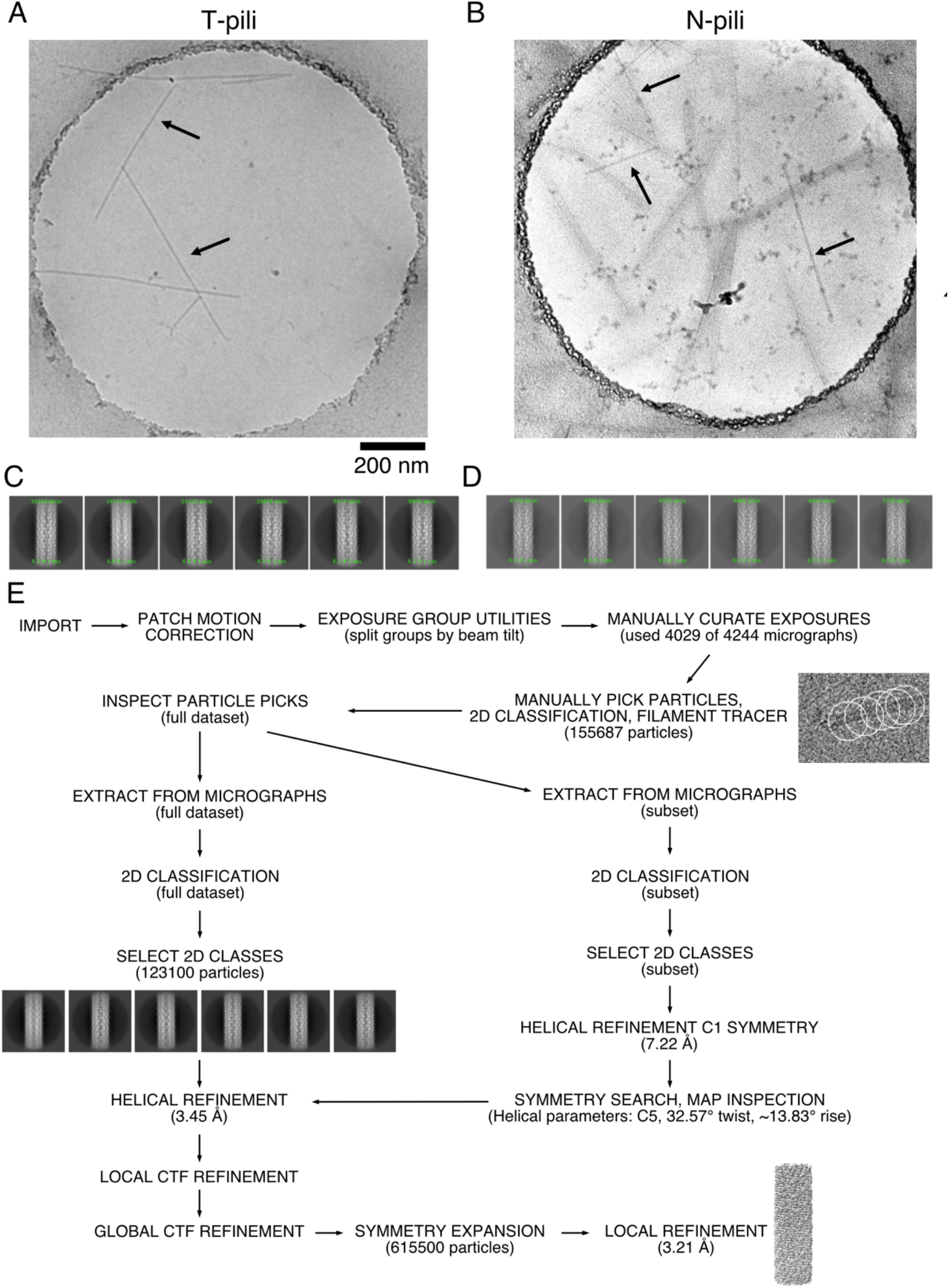
Examples of low-resolution electron micrographs of T-pilus (A) and N-pilus (B). The scale bar represents 200 nm. The difference in the appearance of the micrograph in the N-pilus case is due to the gold coating and graphene oxide layer on the grid. Examples of 2D class averages from T-pilus (C) and N-pilus (D). (E) The workflow for our data analysis of T-pilus using CryoSPARC. N-pilus is analyzed similarly, except the fact that we used the helical parameters from T-pilus for N-pilus analysis right away.

**Supplementary Figure 3.**
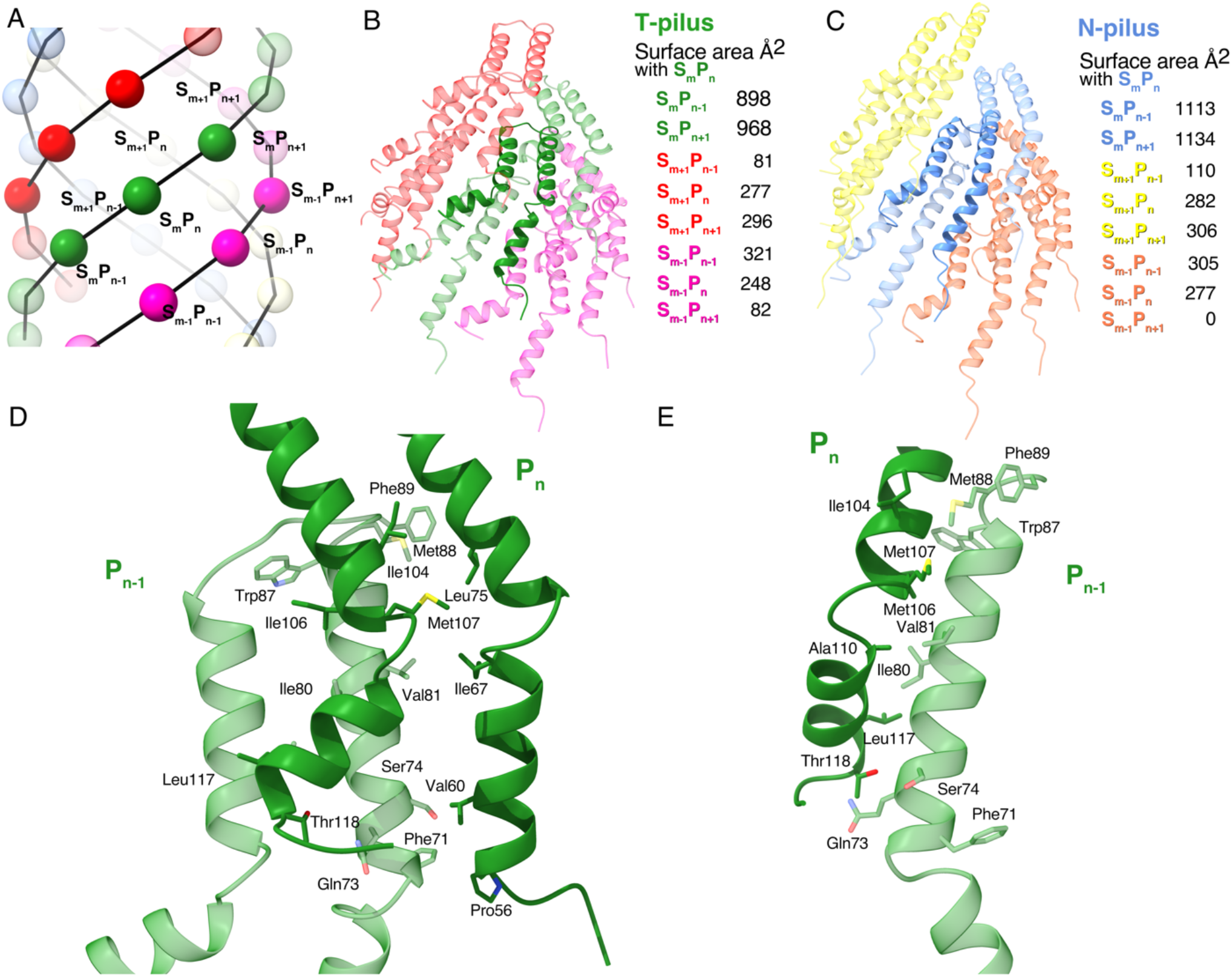
Intermolecular interactions between the different units of VirB2. (A) Schematic cartoon of the T-pilus pilin arrangement showing an interaction unit of nine pilins from three adjacent helical strands. (B) The interaction interface area between other units of VirB2 of T-pilus with the central unit S_m_P_n_ analyzed by PDBsum. The view of the model in (B) is the same as the model in (A). (C) The interaction interface area between other units of TraM of N-pilus with the central unit S_m_P_n_ analyzed by PDBsum. (D, E) The interaction between VirB2 unit n and n-1 analyzed by PDBsum.

**Supplementary Figure 4.**
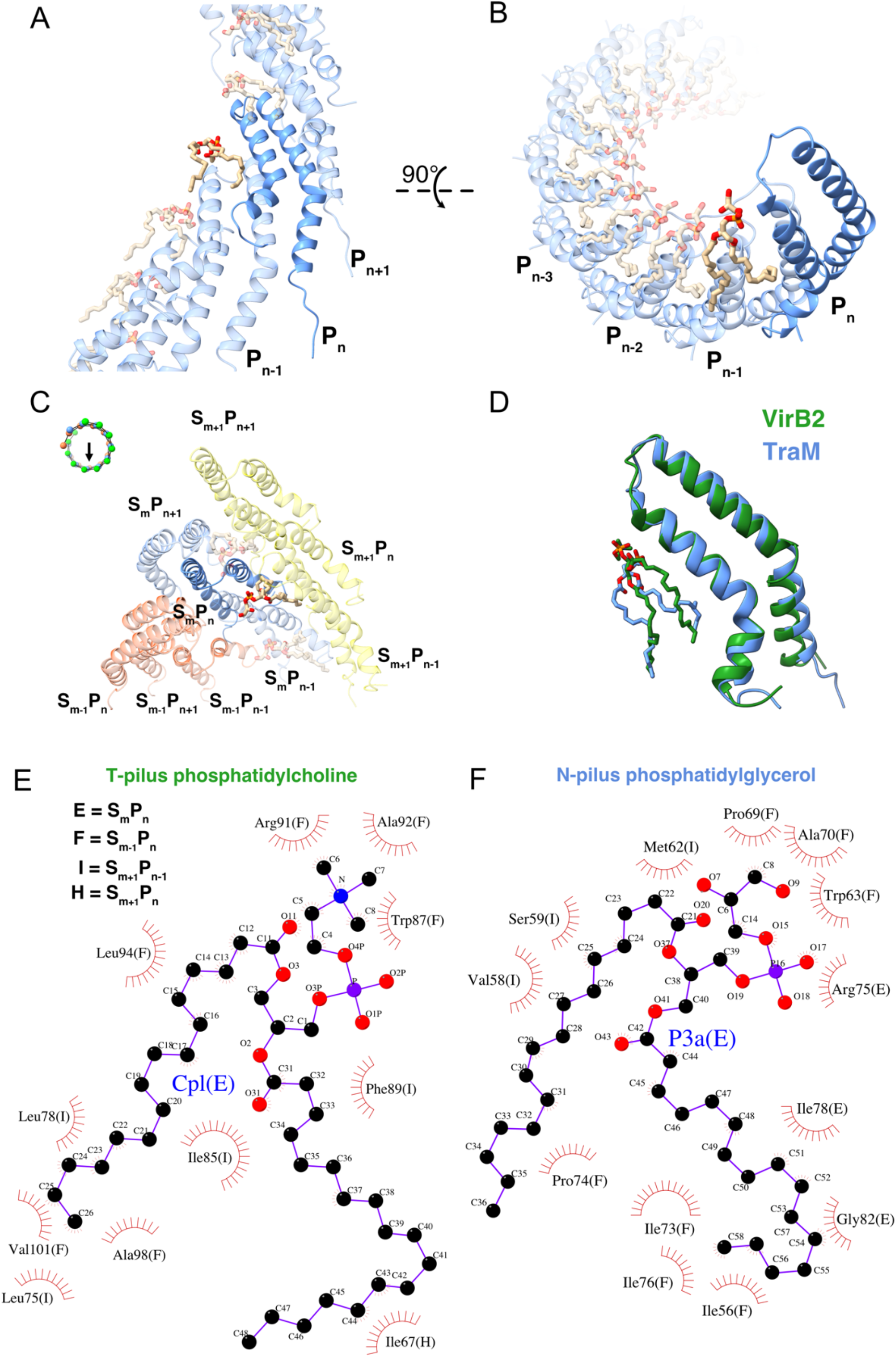
Interactions between phospholipid and pilins. A) Like T-pili (Fig. 3), N-pili contain phospholipid with a stoichiometry of 1:1 with TraM. Arrangement of TraM pilin and lipid in a single helical strand in two different views (A) and (B). (C) Interaction of TraM pilin and phospholipids within a nine-pilin unit similar to Fig. 4D. The small cartoon showing the view in (C) indicated by the black arrow. (D) Overlay of VirB2 and TraM shows that the phospholipid conformation relative to the pilin are significantly different despite the good alignment between VirB2 and TraM. (E) The interactions between phosphatidylcholine and VirB2 within the T-pilus analyzed by PDBsum. (F) The interactions between phosphatidylglycerol and TraM within the N-pilus.

**Supplementary Figure 5.**
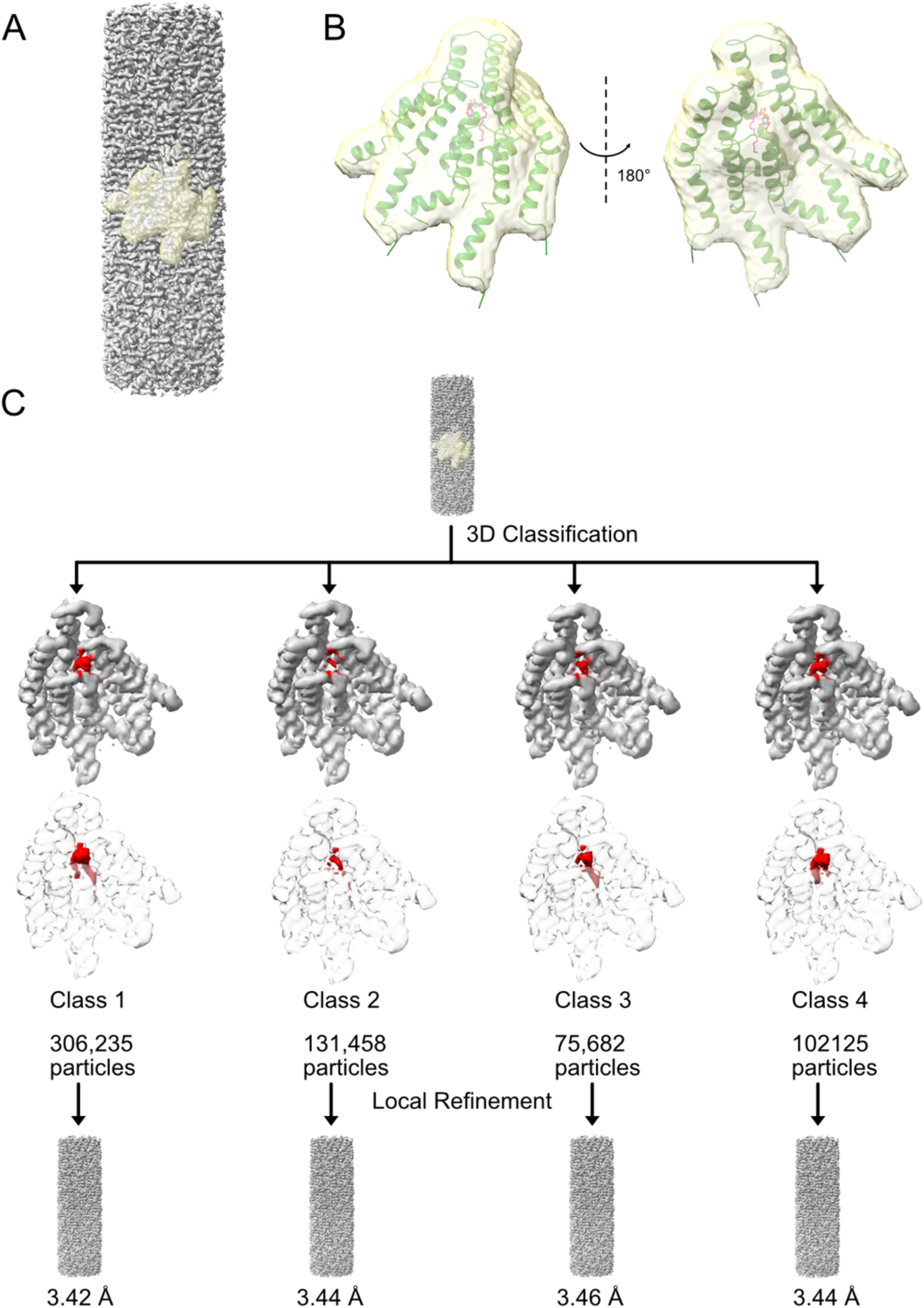
The classification scheme for Lipid Density in one asymmetric unit. (A) The consensus T-pilus 3D reconstruction from 615,500 symmetry expanded particles with the classification mask. (B) The classification mask is overlaid with the corresponding model. The classification mask was designed to cover a single lipid and four surrounding VirB2 molecules. (C) Classification of the symmetry expanded particles into four classes using supervised classification from Cryosparc. The four classes show similar VirB2 density but distinct lipid densities. To avoid model bias, particles of each class were refined with the consensus map. Despite using the consensus map as the model, the resulting maps show clear different lipid densities in the lipid pocket of the asymmetric unit.

**Supplementary Figure 6.**
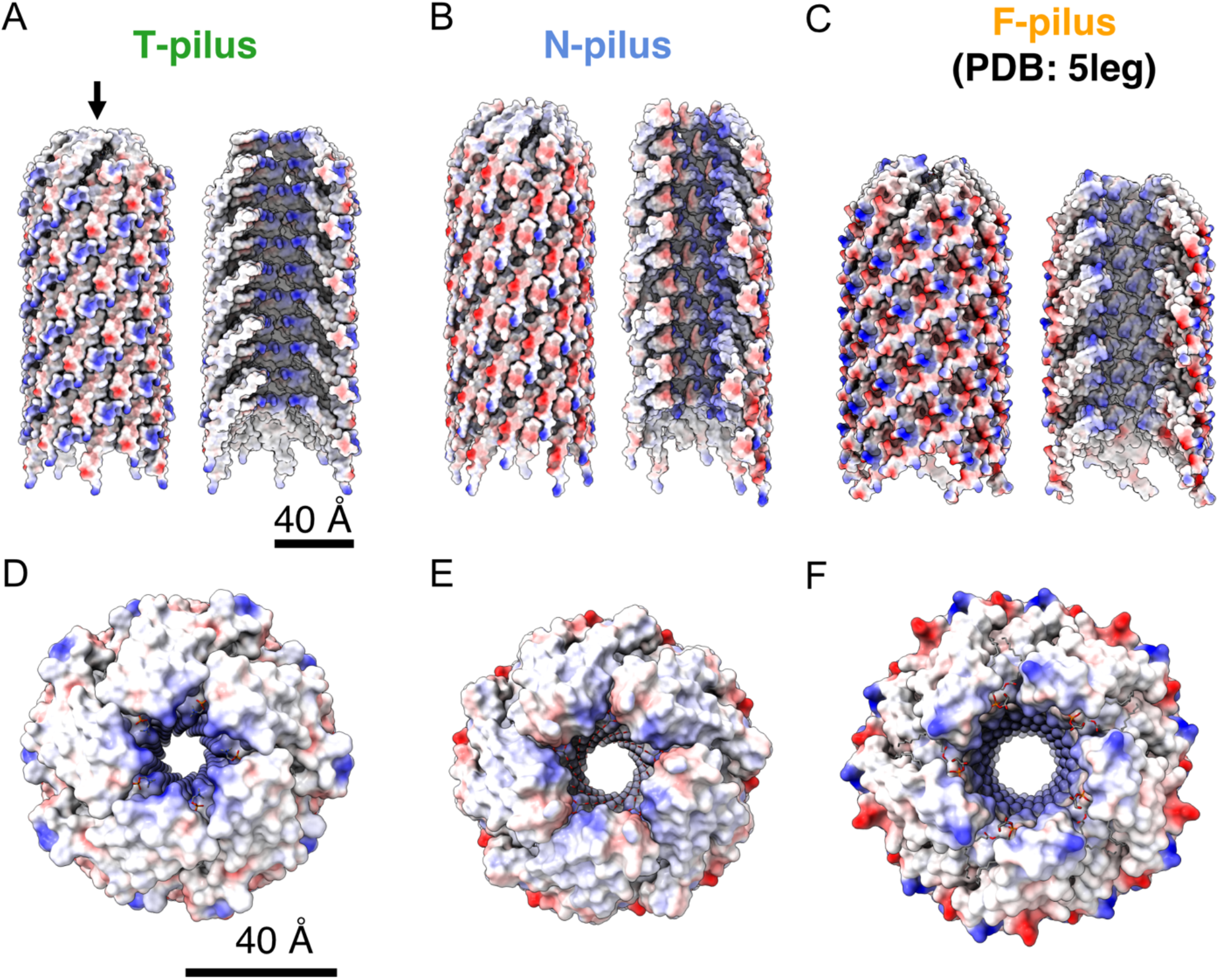
The surface charge analysis of T-pilus, N-pilus and F-pilus. Surface charge analysis of pili only indicates a significant variation in surface charges outside and inside the channel among three types of T4SS pili. (A) Surface charge of the outside of T-pilus (left panel) and inside the lumen by hiding pilins at the front (right side). Surface charges of N-pilus and F-pilus (PDB 5leg) are shown in (B) and (C). The surface charges of the T-pilus (D), N-pilus (E) and F-pilus (F) from top view, looking down the lumen as indicated by the black arrow in (A).

## References

1. H. H. Hwang, M. Yu, E. M. Lai, Agrobacterium-mediated plant transformation: biology and applications. Arabidopsis Book 15, e0186 (2017).

2. K. Dhama et al., Plant-based vaccines and antibodies to combat COVID-19: current status and prospects. Hum Vaccin Immunother 16, 2913–2920 (2020).

3. S. Rosales-Mendoza, V. A. Marquez-Escobar, O. Gonzalez-Ortega, R. Nieto-Gomez, J. I. Arevalo-Villalobos, What Does Plant-Based Vaccine Technology Offer to the Fight against COVID-19? Vaccines (Basel) 8 (2020).

4. P. Vikesland et al., Differential Drivers of Antimicrobial Resistance across the World. Acc Chem Res 52, 916–924 (2019).

5. C. O. Vrancianu, L. I. Popa, C. Bleotu, M. C. Chifiriuc, Targeting Plasmids to Limit Acquisition and Transmission of Antimicrobial Resistance. Front Microbiol 11, 761 (2020).

6. M. Trokter, C. Felisberto-Rodrigues, P. J. Christie, G. Waksman, Recent advances in the structural and molecular biology of type IV secretion systems. Current opinion in structural biology 27C, 16–23 (2014).

7. V. Chandran Darbari, G. Waksman, Structural Biology of Bacterial Type IV Secretion Systems. Annu Rev Biochem 84, 603–629 (2015).

8. T. R. D. Costa et al., Type IV secretion systems: Advances in structure, function, and activation. Molecular microbiology 115, 436–452 (2021).

9. R. Fronzes, P. J. Christie, G. Waksman, The structural biology of type IV secretion systems. Nature reviews. Microbiology 7, 703–714 (2009).

10. R. Fronzes et al., Structure of a type IV secretion system core complex. Science 323, 266–268 (2009).

11. H. H. Low et al., Structure of a type IV secretion system. Nature 508, 550–553 (2014).

12. E.-M. Lai, C. I. Kado, Processed VirB2 is the major subunit of the promiscuous pilus of Agrobacterium tumefaciens. J. Bacteriol. 180, 2711–2717 (1998).

13. K. A. Aly, C. Baron, The VirB5 protein localizes to the T-pilus tips in Agrobacterium tumefaciens. Microbiology 153, 3766–3775 (2007).

14. H. Schmidt-Eisenlohr et al., Vir proteins stabilize VirB5 and mediate its association with the T pilus of Agrobacterium tumefaciens. J. Bacteriol. 181, 7485–7492 (1999).

15. H. Schmidt-Eisenlohr, N. Domke, C. Baron, TraC of IncN plasmid pKM101 associates with membranes and extracellular high molecular weight structures in Escherichia coli. J. Bacteriol. 181, 5563–5571 (1999).

16. T. Kwok et al., Helicobacter exploits integrin for type IV secretion and kinase activation. Nature 449, 862–866 (2007).

17. J. Conradi et al., An RGD helper sequence in CagL of Helicobacter pylori assists in interactions with integrins and injection of CagA. Front Cell Infect Microbiol 2, 70 (2012).

18. B. Hu, P. Khara, P. J. Christie, Structural bases for F plasmid conjugation and F pilus biogenesis in Escherichia coli. Proceedings of the National Academy of Sciences of the United States of America 116, 14222–14227 (2019).

19. K. G. Anthony, C. Sherbourne, R. Sherburne, L. S. Frost, The role of the pilus in recipient cell recognition during bacterial conjugation mediated by F-like plasmids. Mol. Microbiol. 13, 939–953 (1994).

20. R. Eisenbrandt et al., Conjugative pili of IncP plasmids, and the Ti plasmid T pilus are composed of cyclic subunits. J. Biol. Chem. 274, 22548–22555 (1999).

21. P. Khara, L. Song, P. J. Christie, B. Hu, In Situ Visualization of the pKM101-Encoded Type IV Secretion System Reveals a Highly Symmetric ATPase Energy Center. mBio 12, e0246521 (2021).

22. A. L. Samuels, E. Lanka, J. E. Davies, Conjugative junctions in RP4-mediated mating of Escherichia coli. J. Bacteriol. 182, 2709–2715 (2000).

23. M. B. Dürrenberger, W. Villiger, T. Bächi, Conjugational junctions: morphology of specific contacts in conjugating Escherichia coli bacteria. J. Struct. Biol. 107, 146–156 (1991).

24. J. Aguilar, T. A. Cameron, J. Zupan, P. Zambryski, Membrane and core periplasmic Agrobacterium tumefaciens virulence Type IV secretion system components localize to multiple sites around the bacterial perimeter during lateral attachment to plant cells. MBio 2, e00218–00211 (2011).

25. J. Aguilar, J. Zupan, T. A. Cameron, P. C. Zambryski, Agrobacterium type IV secretion system and its substrates form helical arrays around the circumference of virulence-induced cells. Proceedings of the National Academy of Sciences of the United States of America 107, 3758–3763 (2010).

26. H. Y. Wu, C. Y. Chen, E. M. Lai, Expression and functional characterization of the Agrobacterium VirB2 amino acid substitution variants in T-pilus biogenesis, virulence, and transient transformation efficiency. PLoS One 9, e101142 (2014).

27. M. Kalkum, R. Eisenbrandt, R. Lurz, E. Lanka, Tying rings for sex. Trends Microbiol. 10, 382–387 (2002).

28. T. R. D. Costa et al., Structure of the Bacterial Sex F Pilus Reveals an Assembly of a Stoichiometric Protein-Phospholipid Complex. Cell 166, 1436–1444 e1410 (2016).

29. W. Zheng et al., Cryoelectron-Microscopic Structure of the pKpQIL Conjugative Pili from Carbapenem-Resistant Klebsiella pneumoniae. Structure 28, 1321–1328 e1322 (2020).

30. Q. Yuan et al., Identification of the VirB4-VirB8-VirB5-VirB2 pilus assembly sequence of type IV secretion systems. J. Biol. Chem. 280, 26349–26359 (2005).

31. K. A. Aly, L. Krall, F. Lottspeich, C. Baron, The type IV secretion system component VirB5 binds to the trans-zeatin biosynthetic enzyme Tzs and enables its translocation to the cell surface of Agrobacterium tumefaciens. J. Bacteriol 190, 1595–1604 (2007).

32. H. Schägger, G. von Jagow, Tricine-sodium dodecyl sulfate-polyacrylamide gel electrophoresis for the separation of proteins in the range of 1 to 100 kDa. Anal. Biochem. 166, 368–379 (1987).

33. C. Mary, A. Fouillen, B. Bessette, A. Nanci, C. Baron, Interaction via the N-terminus of the type IV secretion system (T4SS) protein VirB6 with VirB10 is required for VirB2 and VirB5 incorporation into T-pili and for T4SS function. J. Biol. Chem. 293, 13415–13426 (2018).

34. J. Snijder et al., Vitrification after multiple rounds of sample application and blotting improves particle density on cryo-electron microscopy grids. J Struct Biol 198, 38–42 (2017).

35. D. N. Mastronarde, Automated electron microscope tomography using robust prediction of specimen movements. J Struct Biol 152, 36–51 (2005).

36. A. Punjani, J. L. Rubinstein, D. J. Fleet, M. A. Brubaker, cryoSPARC: algorithms for rapid unsupervised cryo-EM structure determination. Nature methods 14, 290–296 (2017).

37. D. Liebschner et al., Macromolecular structure determination using X-rays, neutrons and electrons: recent developments in Phenix. Acta Crystallogr D Struct Biol 75, 861–877 (2019).

38. P. Emsley, B. Lohkamp, W. G. Scott, K. Cowtan, Features and development of Coot. Acta crystallographica. Section D, Biological crystallography 66, 486–501 (2010).

39. E. F. Pettersen et al., UCSF Chimera--a visualization system for exploratory research and analysis. J Comput Chem 25, 1605–1612 (2004).

40. E. F. Pettersen et al., UCSF ChimeraX: Structure visualization for researchers, educators, and developers. Protein science : a publication of the Protein Society 30, 70–82 (2021).

41. R. A. Laskowski, J. Jablonska, L. Pravda, R. S. Varekova, J. M. Thornton, PDBsum: Structural summaries of PDB entries. Protein Sci. 27, 129–134 (2018).

42. R. A. Laskowski, M. B. Swindells, LigPlot+: Multiple Ligand-Protein Interaction Diagrams for Drug Discovery. Journal of Chemical Information and Modeling 51, 2778–2786 (2011).

43. A. Forest et al., Comprehensive and Reproducible Untargeted Lipidomic Workflow Using LC-QTOF Validated for Human Plasma Analysis. J Proteome Res 17, 3657–3670 (2018).

44. J. Godzien et al., Rapid and Reliable Identification of Phospholipids for Untargeted Metabolomics with LC-ESI-QTOF-MS/MS. J Proteome Res 14, 3204–3216 (2015).

45. J. Lederberg, E. L. Tatum, Sex in bacteria; genetic studies, 1945-1952. Science 118, 169–175 (1953).

46. J. Manchak, K. G. Anthony, L. S. Frost, Mutational analysis of F-pilin reveals domains for pilus assembly, phage infection and DNA transfer. Molecular microbiology 43, 195–205 (2002).

47. J. E. Kerr, P. J. Christie, Evidence for VirB4-mediated dislocation of membrane-integrated VirB2 pilin during biogenesis of the Agrobacterium VirB/VirD4 type IV secretion system. J Bacteriol 192, 4923–4934 (2010).

48. T. D. Lawley, W. A. Klimke, M. J. Gubbins, L. S. Frost, F factor conjugation is a true type IV secretion system. FEMS Microbiol Lett 224, 1–15 (2003).

49. N. Van Larebeke et al., LARGE PLASMID IN AGROBACTERIUM-TUMEFACIENS ESSENTIAL FOR CROWN GALL-INDUCING ABILITY. Nature 252, 169–170 (1974).

50. H. Schmidt-Eisenlohr et al., Vir proteins stabilize VirB5 and mediate its association with the T pilus of Agrobacterium tumefaciens. J. Bacteriol. 181, 7485–7492 (1999).

